# Role of van der Waals, electrostatic, and hydrogen-bond interactions for the relative stability of cellulose Iβ and II crystals

**DOI:** 10.1101/2024.03.04.583382

**Authors:** Richard Kullmann, Martina Delbianco, Christian Roth, Thomas R. Weikl

## Abstract

Naturally occuring cellulose Iβ with its characteristic parallel orientation of cellulose chains is less stable than cellulose II, in which neighbouring pairs of chains are oriented antiparallel to each other. While the distinct hydrogen-bond patterns of these two cellulose crystal forms are well established, the energetic role of the hydrogen bonds for crystal stability, in comparison to the van der Waals and overall electrostatic interactions in the crystals, is a matter of current debate. In this article, we investigate the relative stability of cellulose Iβ and II in energy minimizations with classical force fields. We find that the larger stability of cellulose II results from clearly stronger electrostatic interchain energies that are only partially compensated by stronger van der Waals interchain energies in cellulose Iβ. In addition, we show that a multipole description of hydrogen bonds that includes the whole COH groups of donor and acceptor oxygen atoms leads to consistent interchain hydrogen-bond energies that account for roughly 70% and 75% of the interchain electrostatics in cellulose Iβ and II, respectively.

## Introduction

Cellulose is the most abundant biopolymer and a sustainable source for a large variety of materials ^1–3^. Naturally occurring cellulose biopolymers are assembled in crystalline arrays, termed cellulose I, in which the polymer chains are oriented in parallel to each other ^4–6^. Cellulose I, however, is not the most stable crystalline assembly of cellulose chains. Dissolving and recrystallizing cellulose I leads to cellulose II ^7^, in which neighboring chains are oriented antiparallel to each other ^8^. For synthetic ^9^ or enzymatically generated ^10^ cellulose oligosaccharides, only cellulose II is observed in crystalline assemblies. Cellulose I and II have characteristic, distinct hydrogen-bond patterns established decades ago ^5,8^(see Fig. 1), but the energetic role of these hydrogen bonds for crystal stability, compared to van der Waals, hydrophobic, and the overall electrostatic interactions, is still a matter of current debate ^11–15^.

**Fig. 1.**
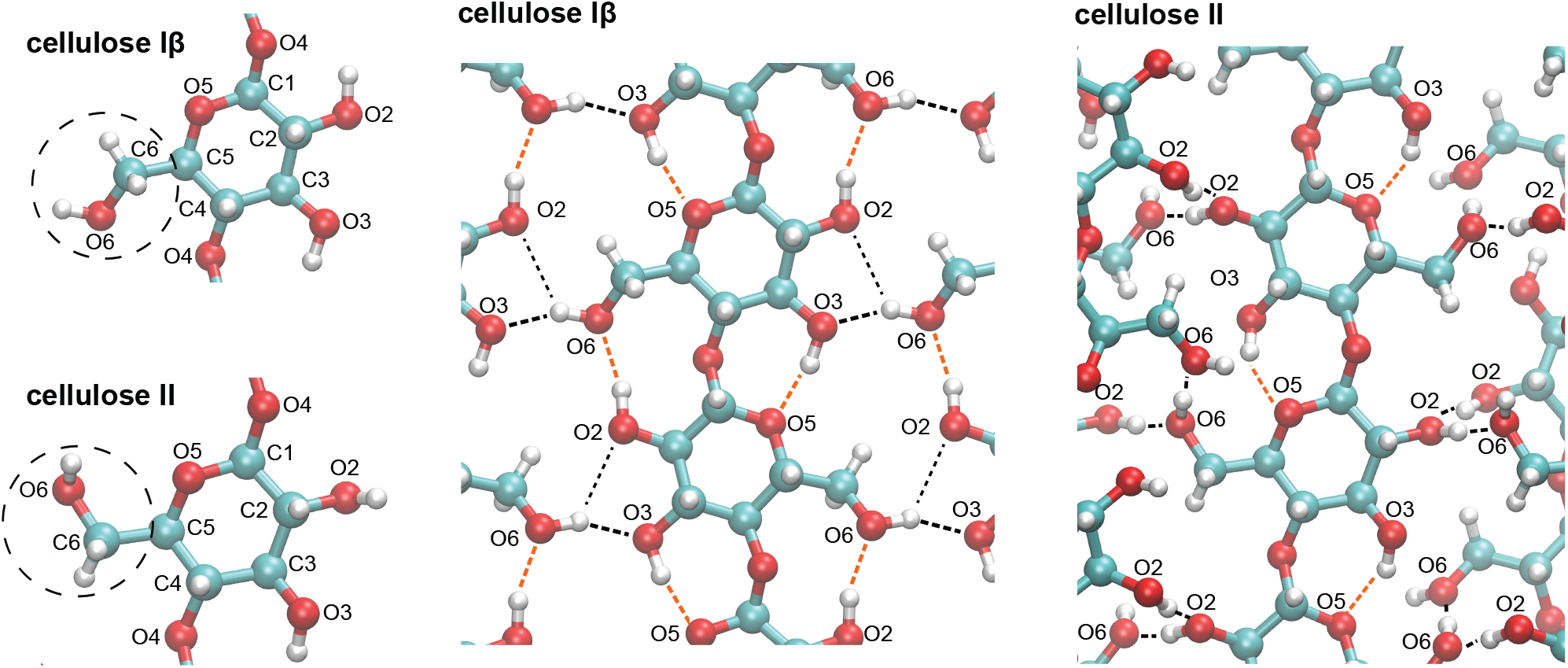
Molecular conformations and hydrogen bonds in cellulose Iβ and II.

Molecular modelling and simulations have been used extensively to explore the hydrogen-bond networks ^16,17^and unit cell parameters ^18^ of cellulose crystals, the twist of cellulose I fibrils ^19–21^, the elastic ^22–26^and thermal response ^27–29^of cellulose, and the assembly and interactions of few cellulose chains ^30–32^. The electrostatic and van der Waals (or London dispersion) intrachain and interchain energies in cellulose crystals have been recently calculated with density functional theory (DFT) methods in conjunction with three popular generations of dispersion corrections ^13^, which lead to differences in dispersion energies of up to about 50% ^13^. The dispersion corrections are necessary to empirically include the long-range dispersion interactions in the approximative quantum-mechanical DFT approach ^33,34^. In classical atomistic force fields, long-range van der Waals interactions are included in the Lennard-Jones pair interaction of atoms (see Methods). The mathematical form and numerous atom-type-specific parameters of force fields have been optimised over decades ^35^, in particular for proteins, resulting in rather accurate descriptions of the structure and dynamics of proteins ^36,37^. Current standard carbohydrate force fields tend to overestimate attractive carbohydrate-carbohydrate interactions in carbohydrate solutions, which has led to recalibrations of the Lennard-Jones potentials for the van der Waals interactions ^38–41^.

In this article, we investigate the relative stability of cellulose Iβ, the dominant form of cellulose I, and cellulose II in energy minimizations with the popular standard force field GLYCAM06^42^ and the recalibrated force field 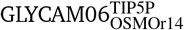 ^39^ starting from the experimentally determined crystal structures ^5,43^. From interpolations of minimization results for different crystal sizes to eliminate surface effects, we determine the ground-state (or “zero-temperature”) bulk energy of the crystal structures in both force fields and obtain a bulk energy difference of several kcal/mol per glucose ring in favour of cellulose II. This bulk energy difference arises from differences in the electrostatic and van der Waals inter-chain energies of cellulose Iβ and II, i.e. from clearly stronger electrostatic interchain energies in cellulose II that are only partially compensated by stronger van der Waals interchain energies in cellulose Iβ. To determine the energetic contributions of the hydrogen bonds formed by three OH groups of the glucose monomers, we propose a multipole description that includes the C atom to which these OH groups are bound as third atom, because the O atoms of the hydroxyl groups “draw” their negative partial charge both from the bound H and C atoms. We show that these multipole description of hydrogen bonds leads to consistent hydrogen-bond energies in good agreement with estimates based on infrared band shifts for cellulose Iβ ^12^.

## Methods

Our energy calculations are based on energy-minimized structures of cellulose Iβ and II nanocystals composed of 52 cellulose 6-mers, 8-mers, 10-mers, and 12-mers. These nanocrystal differ in their volume-to-surface ratio, which we use to extract bulk (volume) energies of cellulose Iβ and II (see Results). We generated initial structures for these energy minimizations from the experimentally determined structures of cellulose Iβ crystals ^5^ and cellulose II crystals ^43^with the software cellulose-builder ^44^, solvated the structures, and performed a first minimization round of 5000 steps of the steepest-descent method followed by 15000 steps of the conjugate-gradient minimization method, in which the cellulose atoms were harmonically restrained with a force constant of 25 kcal mol^*−*1^Å^*−*2^. In second minimization rounds, we fully removed the constraints on cellulose atoms, and generated six energy-minimized structures per crystal by varying the number of the initial minimization steps with the steepest-descent method from 2000 to 7000 in steps of 1000. The minimizations were completed with conjugate-gradient steps to reach a total number of 20000 minimization steps. Our energy calculations include averages over the energy-minimized structures per crystal.

From the partial charges *q* of the atoms in units of the elementary charge, the electrostatic interaction of two atoms *i* and *j* with distance *r* in units of kcal/mol is calculated in the GLYCAM force fields considered here as the Coulomb interaction

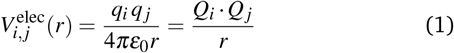

with *Q* = 18.2223 *q*. The partial charges *q* of the atoms in a central glucose ring of the cellulose chains are listed in Table 1. The sum of these partial charges is 0 because the central glucose rings of the cellulose chains are neutral, which leads to overall electrostatic interactions between two such cellulose rings that are short-ranged compared to the Coulomb interactions of atom pairs. The overall electrostatic interactions between two neutral glucose rings are composed of shorter-ranged interactions of charge dipoles and higher charge multipoles. The charged cellulose atoms in Table 1 consist of four groups of atoms that are nearly neutral. To avoid artefacts in the calculation of bulk energies from long-range Coulomb interactions of the charged terminal glucose rings of the cellulose chains, which would be neutralized by the surrounding solvent not considered in our electrostatic calculations, we adjusted the partial charges of the H atoms at the chain termini in these calculations so that also the terminal glucose rings are neutral.

**Table 1.** Partial charges of non-neutral cellulose atoms in GLYCAM force fields and overall charges of the COH groups and remaining atoms with hydrogen–bond-acceptor O5 in units of the elementary charge e.

| atom | charge | group charge |
| --- | --- | --- |
| C2 | 0.310 |  |
| O2 | -0.718 | 0.029 |
| H<sub>O2</sub> | 0.437 |  |
| C3 | 0.284 |  |
| O3 | -0.709 | 0.007 |
| H<sub>O3</sub> | 0.432 |  |
| C6 | 0.282 |  |
| O6 | -0.688 | 0.018 |
| H<sub>O6</sub> | 0.424 |  |
| O4 | -0.468 |  |
| C1 | 0.384 |  |
| O5 | -0.471 | -0.054 |
| C5 | 0.225 |  |
| C4 | 0.276 |  |

The van der Waals interaction is calculated from the Lennard-Jones potential

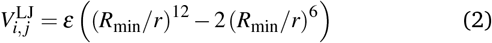

with *R*_min_ = (*R*_*i*_ + *R*_*j*_)*/*2 and 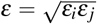 for atom-specific van der Waals radii *R*_*i*_ and *R*_*j*_ and *ε* parameters *ε*_*i*_ and *ε* _*j*_. In the force field 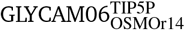, most *ε* parameters of the original force field GLYCAM06 have been slightly rescaled by 0.94 to reproduce experimentally measured osmotic pressures of carbohydrate solutions, which reflect carbohydrate-carbohydrate interactions ^39^. The 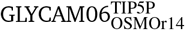 force field employs the TIP5P water model because this water model leads to more reliable carbohydrate-carbohydrate interactions in GLYCAM06, compared to the standard TIP3P water model ^38,45^.

## Results

### Bulk energy of cellulose Iβ and II from minimization

To investigate the relative stability of cellulose Iβ and II in the force fields 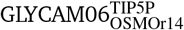 and GLYCAM06, we have analyzed structures of cellulose Iβ and II crystals obtained in energy minimizations with both force fields. Starting structures in these energy minimizations were the experimentally solved structures of cellulose Iβ ^5^ and II ^43^.

The overall energy of a crystal is the sum of its bulk and surface energy. We focus on the bulk energy of cellulose Iβ and II crystals, because the recrystallization of cellulose II from dissolved cellulose Iβ does not seem to be affected by crystal size and therefore likely results from a lower bulk energy of cellulose II compared to cellulose Iβ, and because the surface energies of the crystals include contributions from water interactions that are not directly accessible with energy minimization. To determine the bulk energies of the crystals, we have performed energy minimizations of cellulose Iβ and II crystals composed of 52 cellulose chains with varying numbers of glucose rings per chain. The data points in Fig. 3 represent the interchain electrostatic and van der Waals energy per chain obtained for energy-minimized crystals composed of 52 cellulose 6-mers, 8-mers, 10-mers, and 12-mers. To reduce surface effects from the outer chains in the crystal, the energies in Fig. 3 are averaged over the interchain energies of the 30 central chains in the crystals indicated in Fig. 2. The electrostatic and van der Waals interchain interaction energies of a central chain is calculated as the sum of pairwise energies between the atoms in this chain and the atoms in all other chains of the crystals, divided by two to avoid a double-counting of atom pairs in the averaging over the central chains. The data points in Fig. 3 fall on lines with slopes that reflect energy changes per glucose ring from chain elongation. These energy changes from elongation by glucose rings are equivalent to bulk energies of the cellulose Iβ and II crystals per glucose ring. Table 2 summarizes the interchain electrostatic and van der Waals bulk energies per glucose ring obtained from the linear fits of Fig. 3 with errors estimated as standard errors of the linear fits. The two values per energy term in Table 2 are the energies obtained in the two force fields 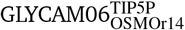(upper value) and GLYCAM06 (lower value). In addition, Table 2 includes the intrachain bulk energy of Iβ and II crystals from linear fitting of the intrachain energies averaged over the 30 central chains of the crystals of 6-mers, 8-mers, 10-mers, and 12-mers akin to Figure 3. The overall bulk energy of cellulose Iβ and II per glucose monomer is the sum of the electrostatic and van der Waals inter-chain energies and the total intrachain energies of Table 2.

**Table 2.** Bulk energies per glucose ring in the force fields 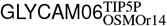 (upper value) and GLYCAM06 (lower value) in kcal/mol.

| | Iβ | II | Iβ - II |
| --- | --- | --- | --- |
| electrostat. interchain | -13.5 ± 0.3<br>-12.4 ± 0.5 | -21.5 ± 0.2<br>-19.8 ± 0.2 | 8.0 ± 0.4<br>7.5 ± 0.5 |
| vdW interchain | -13.0 ± 0.1<br>-15.7 ± 0.1 | -10.1 ± 0.1<br>-12.2 ± 0.1 | -3.0 ± 0.1<br>-3.5 ± 0.1 |
| total intrachain | 115.1 ± 0.3<br>114.6 ± 0.1 | 116.0 ± 0.2<br>116.2 ± 0.3 | -0.9 ± 0.4<br>-1.6 ± 0.6 |
| overall energy | 88.6 ± 0.4<br>86.5 ± 0.5 | 84.4 ± 0.4<br>84.2 ± 0.6 | 4.1 ± 0.6<br>2.3 ± 0.8 |

**Fig. 2.**
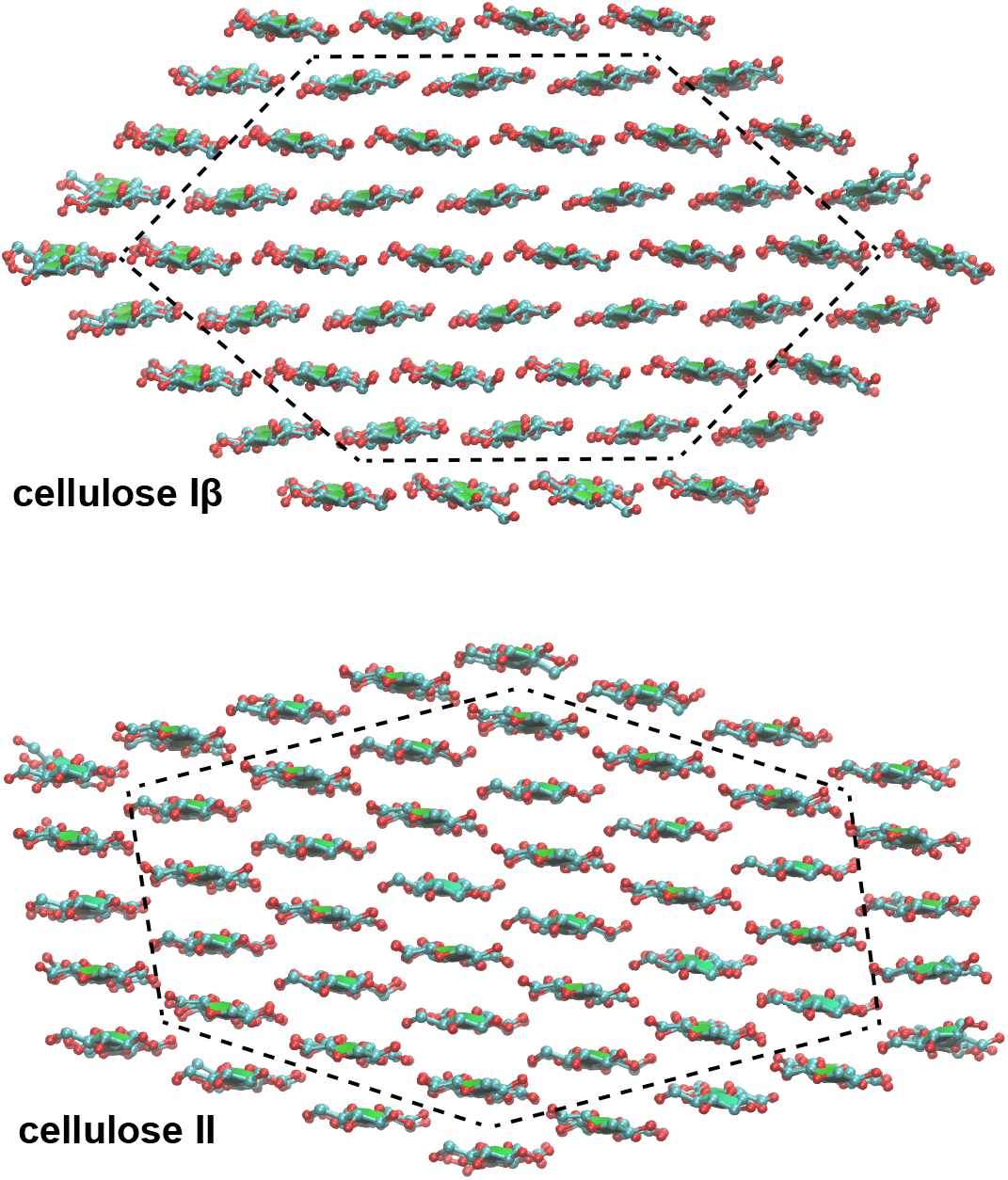
Energy minimized structures of cellulose Iβ and II nanocystals composed of 52 cellulose 6-mers. For clarity, only the four central glucose rings of the 6-mers are shown in the top-view representations of the crystal structures. The dashed lines indicate the 30 central chains of the crystals used in the energy calculations.

**Fig. 3.**
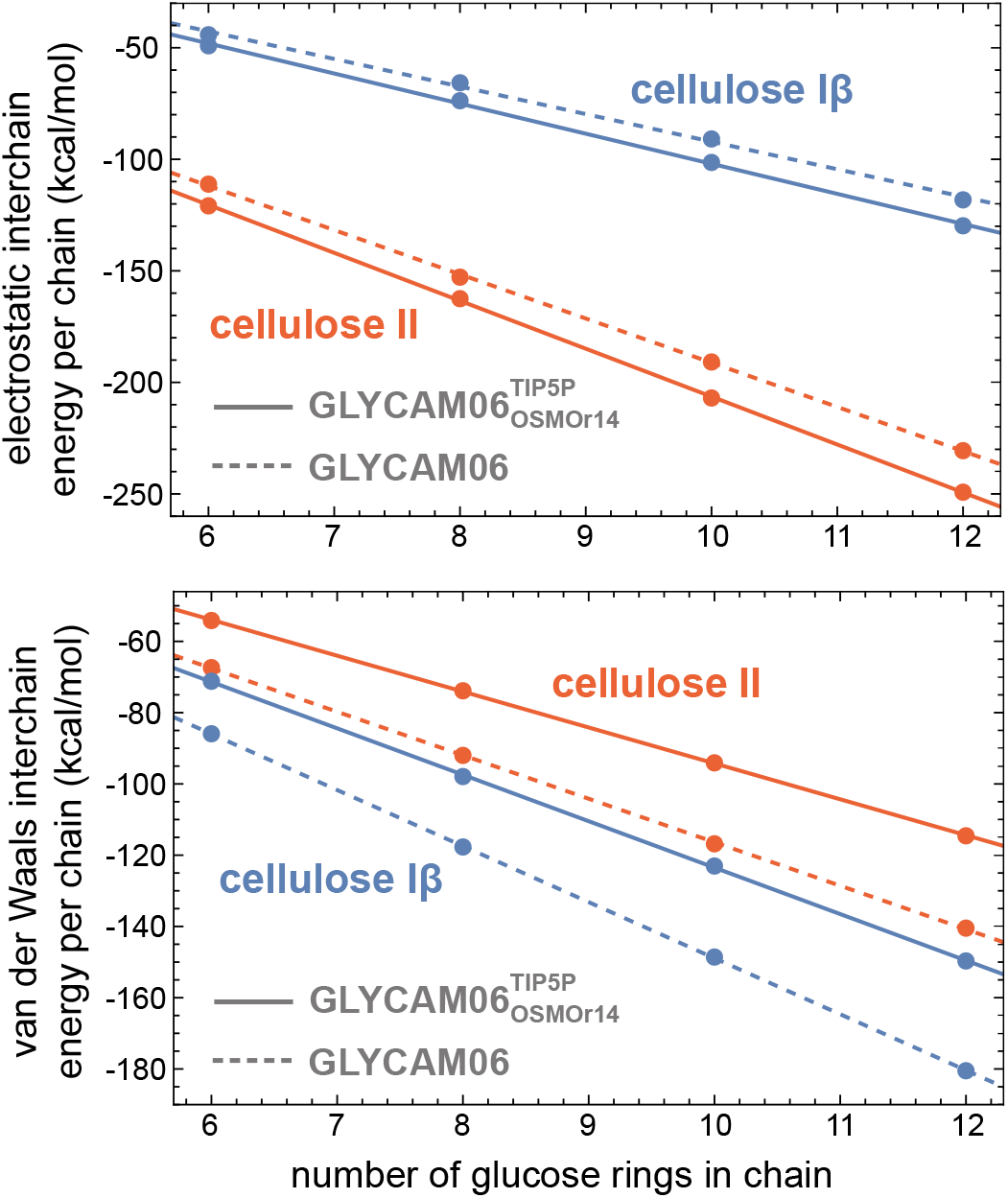
Interpolation of electrostatic and van der Waals interchain energies obtained for the 30 central chains of energy-minimized Iβ and II crystals composed of 52 cellulose 6-mers, 8-mers, 10-mers, and 12 mers (data points). The slope of the fit lines is the bulk interchain energy per glucose ring, i.e. the energy change per glucose ring from chain elongation.

For both force fields, we obtain an overall bulk energy per glucose ring for cellulose II that is several kcal/mol lower than the overall bulk energy for cellulose Iβ (see Table 2). This bulk energy difference arises from differences in the electrostatic and van der Waals interchain energies of cellulose Iβ and II, i.e. from clearly stronger electrostatic interchain energies in cellulose II that are only partially compensated by stronger van der Waals interchain energies in cellulose Iβ. The total intrachain energy in cellulose Iβ and II, in contrast, is rather similar for both force fields, despite the different conformations of the cellulose monomers in both crystals, in particular of the atom O6 of the hydroxymethyl group (see Fig. 1).

### Energies of hydrogen bonds

To assess the role of hydrogen bonds in the interchain interactions, we now focus on the geometry and energies of the hydrogen bonds formed by the three OH groups in cellulose Iβ and II. A simple electrostatic view of hydrogen bonds depicts the OH group of the donor oxygen atom as a dipole with oppositely equal charges *−δ* and +*δ* on the O and H atom, respectively. An electrostatic attraction between the donor OH group and the acceptor O atom with negative partial charge then directly results from the fact that the acceptor O atom is closer to the H atom than to the O atom of the donor group in the hydrogen bond, leading to an attractive Coulomb interaction between H and acceptor O that dominates over the repulsive Coulomb interaction of the two Os. For cellulose, however, the situation is more complex, with an absolute value of the partial charge on the O atom of an OH group in force fields that is significantly larger than the partial charge of the H atom (see Table 1). For the hydrogen bond geometries obtained in our energy-minimized cellulose crystals, the repulsive Coulomb interaction of the two O atoms in a hydrogen bond dominates over the attractive Coulomb interaction between the H atom of the hydrogen bond and the acceptor O atom, leading to an overall positive, repulsive electrostatic interaction between the OH group and the acceptor O. For the exemplary interchain hydrogen-bond O6H-O3 of cellulose Iβ in Table 3, the repulsive electrostatic energy 59.6 kcal/mol between the donor oxygen O6 and acceptor oxygen O3 exceeds the attractive electrostatic energy -56.3 kcal/mol between H_O6_ and O3.

**Table 3.**
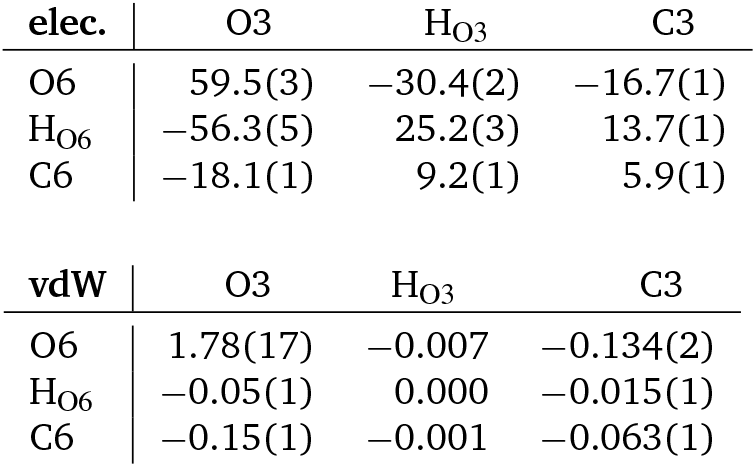
Electrostatic and van der Waals (vdW) interaction energies of H_O6_–O6–C6 and H_O3_–O3–C3 in the intermolecular hydrogen bond O6oH…O3o of cellulose Iβ in the force field GLYCAM (in kcal/mol with standard deviations in brackets)

| elec. | O3 | H<sub>O3</sub> | C3 |
| --- | --- | --- | --- |
| O6 | 59.5(3) | -30.4(2) | -16.7(1) |
| H<sub>O6</sub> | -56.3(5) | 25.2(3) | 13.7(1) |
| C6 | -18.1(1) | 9.2(1) | 5.9(1) |

| vdW | O3 | H<sub>O3</sub> | C3 |
| --- | --- | --- | --- |
| O6 | 1.78(17) | -0.007 | -0.134(2) |
| H<sub>O6</sub> | -0.05(1) | 0.000 | -0.015(1) |
| C6 | -0.15(1) | -0.001 | -0.063(1) |

In the GLYCAM force fields considered here, the negative charge on the O atom of three cellulose OH groups is nearly balanced by the positive charge of the H and C atom bound to the oxygen atom (see Table 1). If we include the atom C6 in the example of Table 3, we obtain a large overall electrostatic attraction of *−*14.9 kcal/mol between the donor group C6-O6-H_O6_ and the acceptor oxygen O3. This large attractive energy helps to understand why the hydrogen bond is formed, but exceeds the overall electrostatic interchain energy of cellulose Iβ per glucose ring (see Table 2). If we also include the C and H atom of the acceptor group, we obtain a total attractive energy of *−*8 kcal/mol between the donor group C6-O6-H_O6_ and the acceptor group C3-O3-H_O3_ as sum of the electrostatic energies in Table 3 between all atoms of the groups. Including all atoms of the nearly neutral donor and acceptor COH groups in the calculation of electrostatic hydrogen-bond energies is reminiscent of the classical approach of Kabsch and Sander ^46^ to determine the energy of hydrogen bonds in protein secondary structures as electrostatic dipole-dipole interactions between the backbone CO group with oppositely equal partial charges *±q*_1_ of the C and O atom and the backbone NH group with oppositely equal partial charges *±q*_2_ of H and N.

Tables 4 and 5 list results for the hydrogen-bond geometry and the electrostatic and van der Waals interaction energies between the donor COH group and acceptor O atom as well as the overall interaction energies between the donor COH group and the acceptor group. Because the crystal cells of cellulose Iβ and II include two chains with slightly different conformations, an origin (o) chain and a center (c) chain ^5,8,12^, we specify and distinguish the hydrogen bonds based on these chain types using a standard distance- and angle-based geometric criterion for identifying hydrogen bonds. The results in Tables 4 and 5 are averages obtained for the hydrogen bonds in the energy-minimized crystals composed of cellulose 12-mers in which the acceptor group is located in the central cellulose chains of the crystals and in the 8 central glucose rings of the 12-mer chains and, thus, in the crystal interior. Numbers in brackets in Tables 4 and 5 indicate standard deviations for the last digit(s) to illustrate variations within the crystal. For hydrogen bonds in which O5 is the acceptor oxygen, we include all 5 atoms indicated in Table 1 as atoms of the acceptor group COX.

**Table 4.**
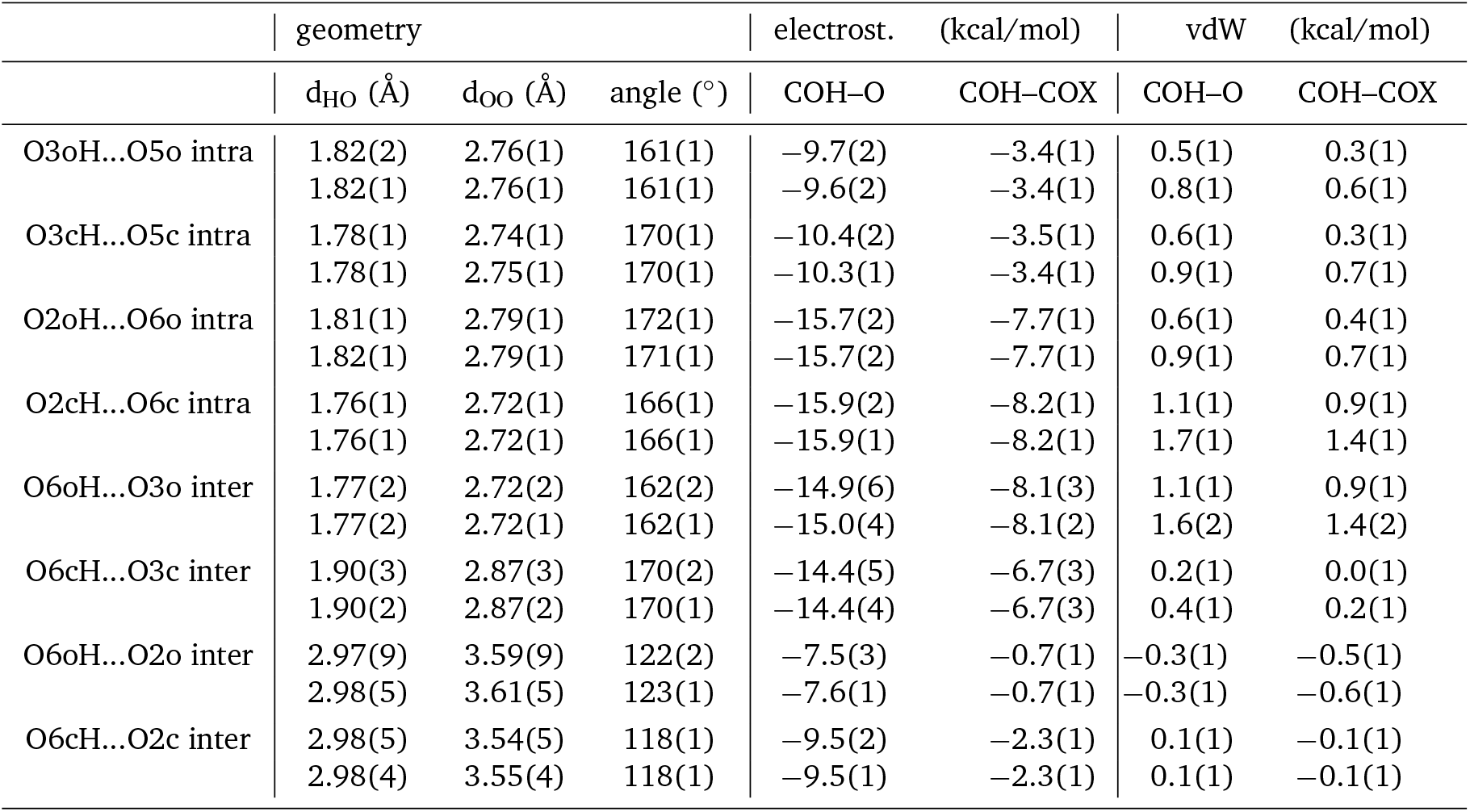
Hydrogen-bond geometry and energetics for cellulose Iβ in the force fields 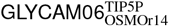 (upper values) and GLYCAM06 (lower values)

|  | geometry |  |  | electrost. (kcal/mol) |  | vdW (kcal/mol) |  |
| --- | --- | --- | --- | --- | --- | --- | --- |
|  | d<sub>HO</sub> (Å) | d<sub>OO</sub> (Å) | angle (°) | COH-O | COH-COX | COH-O | COH-COX |
| O3oH...O5o intra | 1.82(2) | 2.76(1) | 161(1) | -9.7(2) | -3.4(1) | 0.5(1) | 0.3(1) |
|  | 1.82(1) | 2.76(1) | 161(1) | -9.6(2) | -3.4(1) | 0.8(1) | 0.6(1) |
| O3cH...O5c intra | 1.78(1) | 2.74(1) | 170(1) | -10.4(2) | -3.5(1) | 0.6(1) | 0.3(1) |
|  | 1.78(1) | 2.75(1) | 170(1) | -10.3(1) | -3.4(1) | 0.9(1) | 0.7(1) |
| O2oH...O6o intra | 1.81(1) | 2.79(1) | 172(1) | -15.7(2) | -7.7(1) | 0.6(1) | 0.4(1) |
|  | 1.82(1) | 2.79(1) | 171(1) | -15.7(2) | -7.7(1) | 0.9(1) | 0.7(1) |
| O2cH...O6c intra | 1.76(1) | 2.72(1) | 166(1) | -15.9(2) | -8.2(1) | 1.1(1) | 0.9(1) |
|  | 1.76(1) | 2.72(1) | 166(1) | -15.9(1) | -8.2(1) | 1.7(1) | 1.4(1) |
| O6oH...O3o inter | 1.77(2) | 2.72(2) | 162(2) | -14.9(6) | -8.1(3) | 1.1(1) | 0.9(1) |
|  | 1.77(2) | 2.72(1) | 162(1) | -15.0(4) | -8.1(2) | 1.6(2) | 1.4(2) |
| O6cH...O3c inter | 1.90(3) | 2.87(3) | 170(2) | -14.4(5) | -6.7(3) | 0.2(1) | 0.0(1) |
|  | 1.90(2) | 2.87(2) | 170(1) | -14.4(4) | -6.7(3) | 0.4(1) | 0.2(1) |
| O6oH...O2o inter | 2.97(9) | 3.59(9) | 122(2) | -7.5(3) | -0.7(1) | -0.3(1) | -0.5(1) |
|  | 2.98(5) | 3.61(5) | 123(1) | -7.6(1) | -0.7(1) | -0.3(1) | -0.6(1) |
| O6cH...O2c inter | 2.98(5) | 3.54(5) | 118(1) | -9.5(2) | -2.3(1) | 0.1(1) | -0.1(1) |
|  | 2.98(4) | 3.55(4) | 118(1) | -9.5(1) | -2.3(1) | 0.1(1) | -0.1(1) |

**Table 5.** Hydrogen-bond geometry and energetics (in kcal/mol) for cellulose II in the force fields 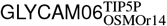 (upper values) and GLYCAM06 (lower values)

|  | geometry |  |  | electrost. (kcal/mol) |  | vdW (kcal/mol) |  |
| --- | --- | --- | --- | --- | --- | --- | --- |
|  | d<sub>HO</sub> (Å) | d<sub>OO</sub> (Å) | angle (°) | COH–O | COH–COX | COH–O | COH–COX |
| O3oH...O5o intra | 1.94(4) | 2.79(2) | 145(3) | −8.5(5) | −3.7(2) | 0.4(1) | 0.3(1) |
|  | 1.93(5) | 2.79(3) | 146(3) | −8.6(6) | −3.7(2) | 0.7(2) | 0.6(2) |
| O3cH...O5c intra | 1.73(2) | 2.68(2) | 164(1) | −10.8(2) | −4.3(2) | 1.1(1) | 0.9(2) |
|  | 1.73(2) | 2.68(2) | 164(1) | −10.8(2) | −4.3(2) | 1.6(2) | 1.5(2) |
| O2cH...O6c inter | 1.71(1) | 2.68(1) | 170(2) | −18.5(4) | −8.6(3) | 1.6(2) | 1.3(2) |
|  | 1.71(2) | 2.68(1) | 170(1) | −18.4(4) | −8.5(3) | 2.2(2) | 2.0(2) |
| O2oH...O2c inter | 1.76(2) | 2.74(2) | 175(2) | −18.5(3) | −7.0(3) | 1.0(2) | 0.8(2) |
|  | 1.75(2) | 2.73(2) | 175(2) | −18.5(4) | −7.0(3) | 1.6(2) | 1.4(2) |
| O6cH...O6o inter | 1.75(2) | 2.73(2) | 173(2) | −16.8(4) | −7.8(2) | 1.1(2) | 0.9(2) |
|  | 1.75(1) | 2.73(2) | 173(2) | −16.8(4) | −7.8(3) | 1.6(2) | 1.4(2) |
| O6oH...O2o inter | 1.73(2) | 2.70(2) | 168(2) | −18.0(4) | −8.1(2) | 1.5(2) | 1.2(2) |
|  | 1.73(1) | 2.70(1) | 168(2) | −17.9(3) | −8.1(3) | 2.1(2) | 1.8(2) |

In cellulose Iβ, there are two intrachain and a branched interchain hydrogen bond per glucose ring. In this branched interchain hydrogen bond, O6 as donor forms a rather strong hydrogen bond with O3 as acceptor and an additional weaker hydrogen bond with O2 as acceptor oxygen. The distances and angles of the hydrogens bonds in Table 4 obtained from our force-field-based energy minimizations are in good agreement with DFT calculations for cellulose Iβ microfibrils ^47^. For the intrachain hydrogen bonds of cellulose Iβ, the distances d_HO_ and d_OO_ obtained from our energy minimizations do not deviate more than 0.05 Å from the distances in the DFT calculations ^47^. For the interchain hydrogen bonds, the distances d_HO_ and d_OO_ of our energy minimizations tend to be slightly larger by about 0.05 Å to 0.2 Å than the distances of the DFT calculations.

In addition to the bond geometries, and in contrast to DFT calculations that allowed only estimates of upper limits of hydrogen bond strengths ^13^, our force-field-based energy minimizations provide detailed insights into the hydrogen-bond energetics. The sum of the electrostatic and van der Waals interaction energies between the COH donor groups and COX acceptor groups in Table 4 ranges from about *−*3 kcal/mol to *−*7 kcal/mol for the intrachain hydrogen bonds and the interchain bonds O6H…O3, in good agreement with the range from *−*4.0 to *−*7.0 kcal/mol of these hydrogen bonds estimated based on infrared band shifts for cellulose Iβ ^12^, which suggests that the hydrogen bond energies in cellulose crystals can be quantified as overall interaction energies of donor and acceptor groups. The electrostatic COH-COH interaction energies of the interchain hydrogen bonds in Table 4 sum up to about *−*8.9 kcal/mol per glucose ring for both force fields, which amounts to 66% and 72% of the overall electrostatic interchain energies per glucose ring in Table 2 for 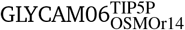 and GLYCAM06, respectively. The electrostatic COH-O interaction energies for the interchain hydrogen bonds, in contrast, sum up to *−*23.2 kcal/mol per glucose ring, which clearly exceeds the overall electrostatic interchain energies per glucose ring in Table 2 and, thus, overestimates the hydrogen-bond energies. The van der Waals interactions of the intrachain hydrogen bonds and the interchain bonds O6H…O3 in Table 4 are repulsive because the distances d_OO_ of the donor and acceptor oxygen atoms in these bonds are clearly smaller than the van der Waals radii 3.442 Å for the oxygen atoms O2, O3, and O6 and the van der Waals radius 3.3674 Å for O5 in the force fields, which leads to positive, repulsive values of the Lennard-Jones potential in Eq. (2).

In cellulose II, the on average two interchain hydrogen bonds per glucose ring occur in different pairings of O2 and O6 as acceptor and donor oxygens in these hydrogen bonds (see Fig. 1 and Table 5). The hydrogen bonds in Table 5 obtained from our energy minimizations correspond to the hydrogen-bond pattern B described by Chen et al. ^16^ as energetically optimal pattern for cellulose II among two alternative patterns. The electrostatic COH-COH interaction energies of the interchain hydrogen bonds in Table 4 sum up to about *−*15.7 to *−*15.8 kcal/mol per glucose ring in the two force fields, which amounts to 73% and 79% of the overall electrostatic interchain energies per glucose ring in Table 2 for 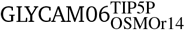 and GLYCAM06, respectively. The electrostatic COH-O interaction energies for the interchain hydrogen bonds, in contrast, sum up to *−*23.2 kcal/mol per glucose ring, which exceeds the overall electrostatic interchain energies per glucose ring in Table 2 and, thus, overestimates the hydrogen-bond energies.

## Discussions and Conclusions

In this article, we demonstrated that energy minimizations with classical force fields can be used to evaluate the energetics of cellulose crystals and of the hydrogen bonds in these crystals. For cellulose Iβ, the atom-atom distances of the hydrogen bonds obtained from our force-field-based energy minimizations in Table 4 are in good agreement with atom-atom distances obtained from DFT calculations for cellulose microfibrils ^47^, and hydrogen-bond energies calculated as the sum of the electrostatic and van der Waals interaction energies between the COH donor groups and COX acceptor groups of the force fields are in good agreement with the hydrogen-bond energies in the range from *−* 4.0 to *−* 7.0 kcal/mol estimated from infrared band shifts for cellulose Iβ ^12^. For cellulose II, the hydrogen-bond pattern obtained from our energy minimizations is the pattern B found to be energetically optimal for cellulose II among two alternative patterns ^16^. As a main result, our force-field-based energy minimizations reproduce the suggested larger stability of cellulose II compared to the native crystal form cellulose Iβ, and trace this larger stability back to clearly stronger electrostatic interchain energies in cellulose II that are only partially compensated by stronger van der Waals interchain energies in cellulose Iβ.

The ranges of the overall electrostatic and van der Waals inter-chain energies per glucose monomer in Table 2 are comparable to ranges recently obtained from density functional theory (DFT) calculations for cellulose crystals ^13^. Depending on the generations of dispersion correction approaches as main error source in the DFT calculations, Li et al. ^13^ obtained values in the range from *−*11.7 to *−*14.8 kcal/mol for the van der Waals interchain energy per glucose monomer in cellulose Iβ, and *−*12.2 to *−*15.3 kcal/mol for the van der Waals interchain energy in cellulose II. From energy minimization in the two forced fields considered here, we obtain the range *−*13.0 to *−*16 kcal/mol for the van der Waals interchain energy in Iβ, and *−* 10 to *−* 12 kcal/mol for cellulose II. For the electrostatic interchain energies per glucose monomer, Li et al. ^13^ obtain the range *−*11.2 to *−*12.4 kcal/mol for cellulose Iβ, and *−*16.7 to *−*17.9 kcal/mol for cellulose II. The ranges of electrostatic interchain energies obtained from our force field minimizations are *−*12.4 to *−*13.5 kcal/mol for cellulose Iβ and *−*19.8 to *−*21.5 kcal/mol per glucose monomer for cellulose II.

For determining the relative stability of cellulose Iβ and II crystals, it is central to note that the dissolved states of the two crystals are identical, and, thus, also the free energies of these states. Stability differences of cellulose Iβ and II crystals therefore need to result from free energy differences of the crystals, and the bulk energies determined from our minimization approach correspond to such free energies in the limit of zero temperature and large crystal size. In principle, stability differences may also result from kinetic rather than thermodynamic free-energy differences, e.g. from different kinetic, or entropic, bottlenecks in the formation or dissolution of two structures. However, at least for cellulose chains composed of rather few glucose monomers, a larger kinetic barrier for forming cellulose Iβ versus II appears implausible.

In summary, we have determined the interchain and intrachain bulk energies in cellulose crystals from linear modeling of force-field-based minimization results for differently sized crystals. Our calculations allow to quantify the role of electrostatic and van der Waals energies in cellulose crystals and provide new insights on the energetics of hydrogen bonds in the crystals. While the dynamics of hydrogen bonds has been well explored in atomistic simulations of aqueous systems ^48,49^, standard approaches focusing on donor OH groups and acceptor O atoms do not lead to realistic descriptions of the hydrogen-bond energetics in cellulose crystals ^12^. We have shown that including the C atoms to which the OH groups are attached in the calculation of hydrogen-bond energies, for both donor and acceptor atom groups, leads to consistent results for hydrogen-bond energies that agree with estimates based on infrared band shifts for cellulose Iβ ^12^.

## Acknowledgements

This work was funded by the International Max Planck Research School (IMPRS) on Multiscale Bio-Systems and the Max Planck Society.

## Availability of data

The structures and force-field terms of the energy-minimized cellulose crystals of this work are available in the Edmond data repository at https://doi.org/10.17617/3.1FPW2C ^50^.

